# Fixed Players, Fluid Moves: Refining Strategy While Preserving Identity in Free-Ranging Dogs

**DOI:** 10.64898/2026.09.15.751833

**Authors:** Tuhin Subhra Pal, Sagarika Biswas, Anindita Bhadra

## Abstract

Animals often display active decision-making while navigating noisy habitats, for varied purposes like foraging, finding shelter, evading predators or finding suitable mates. While some responses are hardwired, following fixed action patterns, others are more flexible, and display context-dependent plasticity. Animals might weigh behavioural consistency against adaptive plasticity when facing recurrent ecological difficulties, but the nuanced mechanisms orchestrating this balance is not well understood. We investigated this question by repeatedly exposing 47 free-ranging dogs to chicken contaminated with lemon solution, an aversive but edible and familiar food source, across four consecutive trials. By integrating behavioural frequencies, latency measures, feeding efficiency, interaction-time and location effects, repeatability analyses, and behavioural sequence structure, we examined how repeated experience influenced foraging behaviour. Contrary to expectations of broad behavioural reorganisation, most behaviours remained stable. Initial approach latency was strongly repeatable within individuals with no detectable influence of sex or trial number across the population. This suggests that how an animal first assesses risk is primarily shaped by stable personality traits rather than momentary external factors. Eating behaviour and bowl nudging also changed selectively across exposures, with modest sex differences in several behaviours. Behavioural sequences became increasingly similar over time, while several behaviours retained consistent individual differences, indicating persistent behavioural tendencies. Our results collectively demonstrated that repeated aversive exposure selectively modified a few critical decision points rather than the entire behavioural repertoire. This implies that animals can adapt by making small adjustments to specific behaviours while maintaining the stability of their overall behavioural architectures. Animals seem to assess risk separately from how well they use resources. That separation may help explain how stable personalities can coexist with the fast, flexible decisions animals need in unpredictable environments shaped by humans.

## Introduction

Animals often repeat specific behaviours, acting in ways that differ by context, from signalling location or arousal to assessing quality in mating or conflict scenarios. Such repetition, though energy-demanding, may reaffirm, replace, or amplify previous signals (Payne & Pagel, 1997), prompting important questions about its adaptive benefit, especially in species navigating complex and variable environments. In dynamic and uncertain environments, animals must weigh behavioural flexibility with routine decision-making to maximize foraging success. While learning enables adaptation to varying conditions (Hughes et al., 1992), cognitive rigidity and trait-driven heuristics can reduce uncertainty and exploration costs in predictable settings (Dias et al., 1996; Balleine & Dickinson, 1998). This aligns with classic ideas in optimal foraging theory: animals are willing to put up with certain costs or risks if the food is good enough (Sih & Christensen, 2001; Sih & Bell, 2008), a pattern recently observed in free-ranging urban dogs who dynamically adjust their sequence of searching, assessing, and resource handling depending on risk and quality of patches (Sarkar et al., 2023). During foraging, they generally favour speed over accuracy, though they sniff out of all options when the payoff is worth it (Sarkar et al., 2025b).

Individual differences in foraging behaviour are well documented across taxa. Bold individuals may tolerate aversive cues to secure high-value rewards, whereas shy counterparts prioritize safety (Sih & Del Giudice, 2012). Such traits can persist as stable behavioural syndromes, as seen in pigeons’ reversal-learning strategies, where individuals consistently act as “anticipators” or “perseverators” (Stagner et al., 2012). Personality-driven consistency has also been demonstrated broadly across vertebrates (Gosling, 2001; Sih et al., 2004). Defined as stable behavioural variation among individuals facing comparable situations (Réale et al., 2007), such traits are reflected in consistent intra-situational behaviours in dogs (Svartberg et al., 2005), pumpkinseed sunfish (Coleman & Wilson, 1998), and dairy goats (Lyons et al., 1988). These stable differences aren’t accidents. They reflect persistent traits (Zuckerman, 1991). But consistency need not be uniform. Proactive–reactive coping styles and exploration–avoidance tendencies are increasingly treated as distinct behavioural axes, not a single trait dimension (Réale et al., 2007; Sih et al., 2004). At least partly independent mechanisms may regulate approach and handling of a risky resource (Sih & Bell, 2008). Experimental evidence shows a continuum from flexibility to rigidity. Even after repeated exposure, zebrafish exhibit the same level of boldness or shyness (Budaev and Brown, 2011). In maze difficulties, rats and mice apply different amounts of space-based tactics or repeat old strategies (Tolman, Ritchie, & Kalish, 1946). Even when the rules of reward change, pigeons continue to provide the previously reinforced response (Stagner et al., 2012). Even when other locations are just as good, honeybees firmly stick to one location (Fragoso and Brunet, 2023). In contrast, rhesus macaques adjust their strategies in response to shifting opportunities for reward (Sugrue et al., 2004). When taken as a whole, this indicates that stable behaviour and flexible behaviour are on the same continuum. Together, these findings imply that strategic consistency and flexibility coexist within a spectrum shaped by species-specific ecology and individual temperament.

Here, associative learning is essential as it aids in developing enduring foraging techniques. By associating sensory signals with post-ingestive findings, animals develop stable dietary preferences or aversions (Mehiel & Bolles, 1988; Sclafani, 1995). The initial hedonic value of a substance is not necessary for nutritional conditioning to take place. Post-ingestive feedback is highlighted by this (Mehiel & Bolles, 1988). Long-lasting aversions can result from delayed negative feedback as well (Burritt & Provenza, 1991). This encourages nutritional diversity and long-term strategic consistency (Burritt & Provenza, 1991; Provenza, 1996). Stable behavioural patterns can be promoted by mild but persistent aversions (Provenza, 1996). Additionally, based on trustworthy post-ingestive feedback, foragers may continuously adjust their preferences, updating the projected net benefits (Sclafani, 1995), with social context sometimes outweighing food value, as when affiliative touch is preferred over food (Nandi et al., 2026). Consistency is also shaped by resource spacing. Repetitive contacts include a trade-off between exploration and exploitation beyond patch residence times (Stephens et al., 2008). Animals strategically sample to acquire information about the reliability of resources. Foragers reduce uncertainty by integrating reinforcement history through Bayesian updating. Once the cost of a resource, like acidity, is predictably determined, they switch from flexible sampling to stable choice based rules (Valone, T.J., 2006). Sensory input and reinforcement history are integrated into stable decision-making rules by neural mechanisms, including as value-encoding processes in the parietal cortex (Sugrue et al., 2004). Delayed aversion learning can further consolidate avoidance strategies (Burritt & Provenza, 1991). But behavioural change across repeated exposures need not always reflect associative learning of this kind. It may equally arise from non-associative, perceptual habituation to the aversive stimulus. Disentangling the two remains a persistent challenge in repeated-exposure paradigms. Thus, consistent foraging behaviour may emerge from interactions among spatial cognition, reinforcement history, and ecological pressures.

These dynamics matter for free-ranging dogs (*Canis lupus familiaris*). They forage in human-dominated landscapes characterized by unpredictability and risk, recognizing individual humans (Sarkar et al., 2024; Sarkar and Bhadra, 2022; Bhattacharjee et al., 2017). They also adapt begging to social cues (Biswas et al., 2026) and read human behavioural cues to guide foraging (Sarkar et al., 2025a). While scavenging from garbage, FRDs frequently encounter non-preferred and inedible items mixed with preferred food sources (Sarkar et al., 2019; Butler et al., 2018). Lemon is common in Indian refuse due to widespread culinary use. This raises the likelihood of recurrent exposure. Human provisioning is essential for pet dogs (Bhattacharjee et al., 2017) and previous studies depicted that they frequently exhibit an aversion to citrus flavors like lemon (Bradshaw, 2006). This aversion most likely reflects an old vertebrate adaptation for identifying acidity in the environment. Later, it was appropriated to identify food that was spoiled or contaminated (Frank et al., 2022). Thus, avoiding sour foods is functionally adaptive. However, FRDs are subject to ecological limitations, unlike domestic dogs. It might not always be possible to reject tainted food (Bhattacharjee et al., 2017; Woodford, 2012; Pal et al., 2025a). This ecological conflict is supported by empirical data. Lemon juice and whole lemons are clearly avoided by adult FRDs (Pal, 2022; Pal et al., 2025a). Lemon pulp and rind are less avoided than lemon juice, and adults preferentially scavenge from less concentrated lemon juice environments over more concentrated ones (Pal et al., 2025a). Recent findings further demonstrate active avoidance of high lemon concentrations and adaptive feeding adjustments based on sensory properties (Pal et al., 2025a). Adults exhibit greater selectivity and deliberate foraging, whereas juveniles display less discernment, highlighting developmental influences (Pal et al., 2025a; Pal et al., 2026). Additionally, free-ranging dogs exhibit sixteen distinct strategic behaviours when scavenging in acidic environments, with high lemon concentrations reducing foraging ability and strategic selection due to aversion (Pal et al., 2025b). Together, these findings indicate both flexibility and constraint in acidic foraging contexts, but whether individual dogs consistently rely on particular strategies across repeated encounters remains unresolved.

Building on these findings (Pal et al., 2025a, b; Pal et al., 2026), we aimed to quantify intra-individual variation (IIV) to understand whether free-ranging dogs maintain stable strategic tendencies or selectively modify behavioural responses across repeated aversive foraging exposures. Specifically, we examined sequence organization to understand whether individuals exhibit consistent behavioural profiles and sequence organization across trials. By assessing whether dogs maintain a strategic fingerprint across trials, we can distinguish between transient behavioural noise and enduring individual dispositions. We hypothesize three, non-mutually exclusive possibilities. If personality traits and learned heuristics dominate decision-making, individual dogs should show consistent strategic profiles across trials. If cognitive flexibility and real-time updating prevail, dogs should progressively and broadly modify their strategies. Alternatively, consistent with an optimal-foraging view in which risk assessment and resource handling are at least partially separable (Sih & Bell, 2008), dogs may preserve stable overall dispositions while selectively refining only those decision points that determine efficiency.

## Materials and methods

### Study Sites

The experiment was carried out in 6 broad areas subdivided into 18 sampling points in the Nadia district (22.9747° N, 88.4337° E) of West Bengal (Supplementary figure 1). Locations were not repeated over the course of data collection to avoid accidental resampling. The experiments were conducted in two time slots: 06:00 -12:00 hrs and 15:00 - 20:00 hrs to avoid unnecessary extraneous human interactions in terms of feeding leftovers from households around typical lunch hours. Flux data were video recorded to quantify human movement and assess its relation to fluctuations in dog behaviour (Bhattacharjee et al., 2021).

### Selection and Identification of Subjects

Only adult free-ranging dogs were included in the experiment (N=47), with adulthood determined based on body size and visible genital structures. All trials were conducted on dogs that appeared healthy, showing no obvious signs of disease or injury, and that voluntarily participated in the experiment. Dogs that exhibited signs of extreme anxiousness, fear, or avoidance behaviour, such as running away during the placement of the experimental setup, were excluded from the study to minimise the influence of prior trauma or stress on behavioural responses. The sex of each dog was recorded during the experiment by direct observation of genitalia. Random sampling was employed, with the primary focus on solitary adult dogs. In addition, dogs that were initially observed in groups of two or three individuals were also included, with one focal individual tested individually per trial. In such cases, one focal individual was selected for the trial, and observations were recorded to assess the potential influence of group presence on the behaviour of the focal dog.

### Materials

We used fresh boneless chicken pieces (15g each), 25% lemon juice solution, biodegradable white paper bowls (capacity 150 ml), and a mobile phone camera for the experiment. All materials, including biodegradable bowls, chicken pieces, and lemon juice solutions were replaced after each trial to avoid contamination and maintain experimental consistency. For each trial, 20 ml of 25% lemon juice solution was poured into a biodegradable bowl, and a 15g chicken piece was placed in the bowl of solution. The prepared bowl was positioned approximately 1–1.5 m in front of the focal dog to allow free interaction with the setup.

## Methodology

### Method of Experiment

Each individual dog participated in four consecutive trials, each lasting 90 seconds, with a minimum inter-trial interval of 30 seconds and a maximum of 60 seconds. One experimenter recorded the trial while the other placed the bowl and then stepped back to a distance of 1.5–2 metres, standing with arms relaxed at the sides and avoiding eye contact with the focal dog (Figure 1). The setup was positioned directly in front of the dog to allow free approach. Bowls were placed on relatively flat surfaces to prevent tilting and to avoid giving the dogs a mechanical advantage in extracting the chicken without contacting the solution. To minimise interference from oncoming traffic, trials were conducted at the side of the road. The entire process was recorded using a handheld mobile camera which was kept at a distance of at least 2 metres from the focal dog (Supplementary video 1).

**Figure 1.**
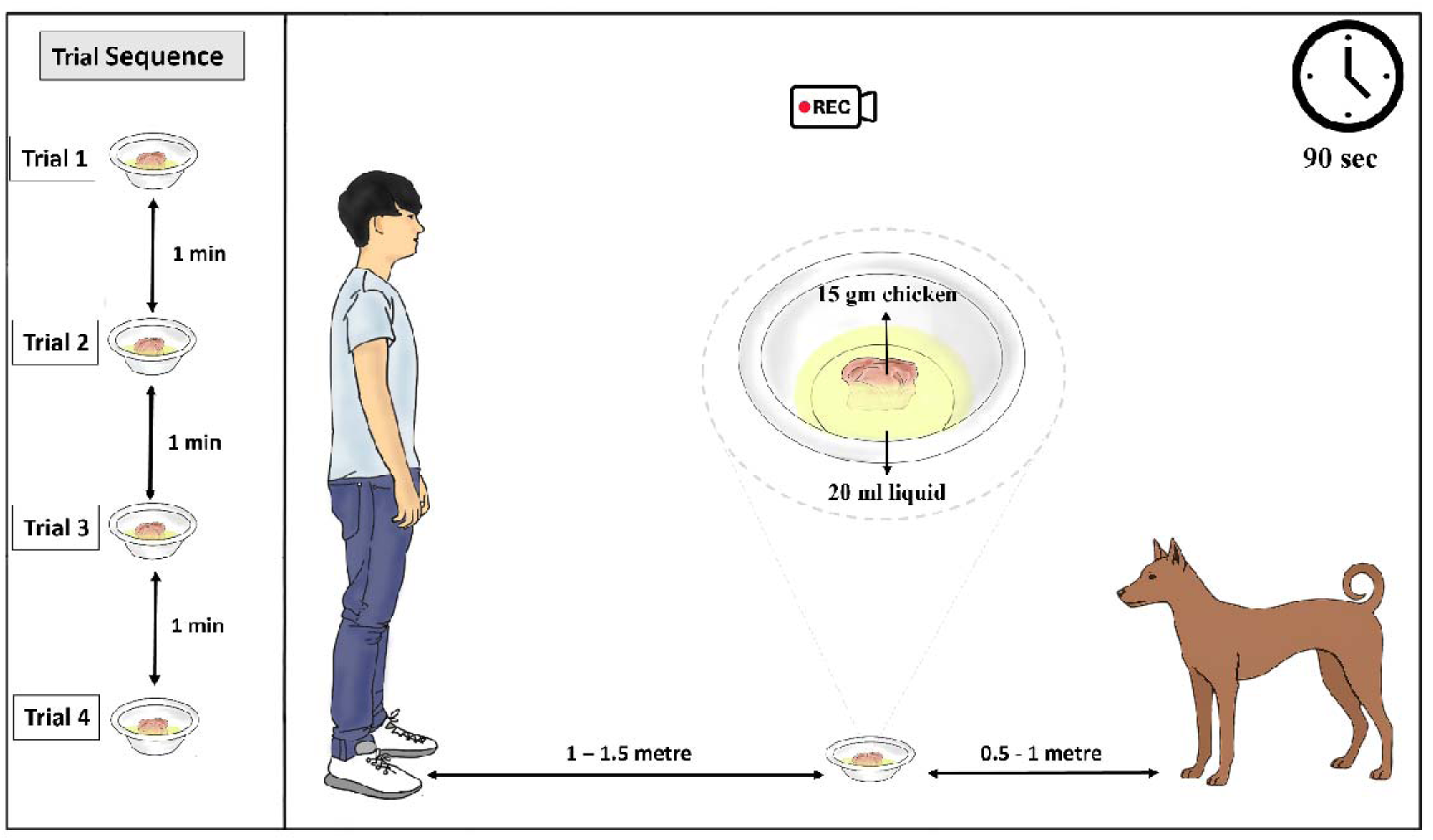
Schematic representation of the experimental set-up.

### Flux Analysis

For each site where an experiment was conducted, flux was calculated as a measure of the degree of human interaction (Bhattacharjee & Bhadra, 2020). Three videos each of one minute were recorded keeping the frame fixed. An imaginary vertical line was ascertained at a central point. The flux was calculated for each video by counting the total number of individuals and vehicles crossing the vertical line (every single vehicle irrespective of the number of pillion riders was counted as one) and then averaged to have a flux value for each dog.

### Behavioural analysis

Videos were decoded for the behavioural analysis. All interactions of the focal dog with the food bowl were recorded from the moment the bowl was placed until completion of eating or termination of engagement. Each behavioural event was documented with its sequence, duration, and contribution to the dog’s overall foraging strategy. Behaviours were classified into predefined categories such as sniffing, licking, eating, food manipulation, and strategic food-placement modifications, as detailed in the ethogram (Supplementary table 1).

Manipulative behaviours such as nudging or upturning the bowl, using forelegs, carrying food away, rubbing, shaking the food, and unsuccessful attempts (e.g., missed grabs) were recorded alongside direct consumption behaviours. Behavioural data were further analysed across experimental conditions, time of day, and location to assess external influences on strategy use. Total interaction time was defined as the cumulative duration of all physical or investigatory interactions with the experimental setup, measured from the first interaction with the bowl, food, or lemon-contaminated solution until the dog stopped engaging with the setup or completed food consumption. This measure did not include the latency to approach the setup. Success in food consumption, number of attempts observed, and total duration from initial sniffing to final consumption were quantified to assess efficiency. Post-consumption behaviours were also recorded to provide a comprehensive evaluation of the animals’ immediate response after consuming the food. Behavioural durations were measured in milliseconds to allow precise quantification of transitions.

## Statistical analysis

All statistical analyses were conducted in R Studio (v4.2.2; R Core Team, 2022). Prior to model fitting, the distribution of continuous response variables was evaluated using the *fitdistrplus* package. Model selection was determined by comparing Akaike Information Criterion (AIC) and Bayesian Information Criterion (BIC) values (Akaike, 1974; Schwarz, 1978). A linear mixed-effects model was used to analyse log-transformed approach latency as a function of trial number and sex. Time investment data were transformed into proportions by dividing the duration of each behavioural category by the total interaction time, and analysed using independent beta regression models via the *betareg* package with trial number and sex as predictors. (Partial eating (PE) and not-visible (NV) instances were excluded from formal statistical modelling due to their low frequency of occurrence across trials, which precluded reliable model convergence.)

Kaplan–Meier survival analyses were performed using the *survival* package, and a mixed-effects Cox proportional hazards model with dog identity as a random intercept was fitted using the *coxme* package to analyse the duration from first sniff to complete food consumption. Trials in which dogs did not complete food consumption within the 90-s observation window were treated as right-censored observations. Adjusted survival curves were plotted using the *survminer* package. Gamma generalized linear mixed models (GLMMs) with a log link and dog identity as a random intercept were fitted using the glmmTMB package to analyze total interaction time (as a function of initial approach latency and location) and sniff-to-lick decision-making latency (as a function of trial number and location). Behavioural repeatability coefficients (R) were estimated with dog identity as a random effect using the *rptR* (Stoffel et al., 2017) and *lme4* packages, supported by data manipulation via *dplyr, tidyr and e1071*. Behavioural sequence analysis was conducted using length-normalized Optimal Matching (OM) distances based on transition-rate substitution costs using the *TraMineR* package (Gabadinho et al., 2011). We used *effects, ggplot2 and reshape2* packages for visualizing and processing model effects, sequence data, and transitions.

## Results

### 1. Does repeated exposure influence latency to approach the aversive food setup?

We evaluated whether repeated exposure influenced the approach latency of free-ranging dogs by fitting a linear mixed-effects model to analyse log-transformed approach latency, with trial number and sex as fixed effects and dog identity fitted as a random intercept.

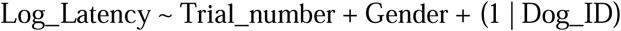

Approach latency remained remarkably consistent across repeated exposures. Dogs approached the aversive food setup with comparable latencies throughout the four trials, indicating a stable behavioural response over time. Relative to the first trial (T1), no significant differences were detected in T2 (β = 0.15, *p* = 0.334), T3 (β = −0.16, *p* = 0.282), or T4 (β = −0.004, *p* = 0.980). Likewise, males and females exhibited similar approach latencies (β = −0.08, *p* = 0.749). Allowing trial-specific random slopes for individual dogs did not significantly improve model fit (likelihood-ratio test: χ² = 13.81, df = 9, *p* = 0.129), indicating no evidence that dogs differed in their behavioural responses across repeated trials.

Approach latency showed significant repeatability among individuals (R = 0.531 ± 0.074, 95% CI = 0.362–0.651, likelihood-ratio test: *p* < 0.001; permutation test: *p* = 0.001), indicating that approximately 53% of the variation in approach latency was attributable to consistent differences among individual dogs (Figure 2). It suggests that behavioural responses were highly consistent within individuals while differing across them, where such variation was independent of gender. Overall, these findings demonstrate that repeated exposure did not alter dogs’ willingness to approach the aversive food setup, highlighting the robustness and repeatability of individual approach behaviour across successive trials (Supplementary information 1).

**Figure 2.**
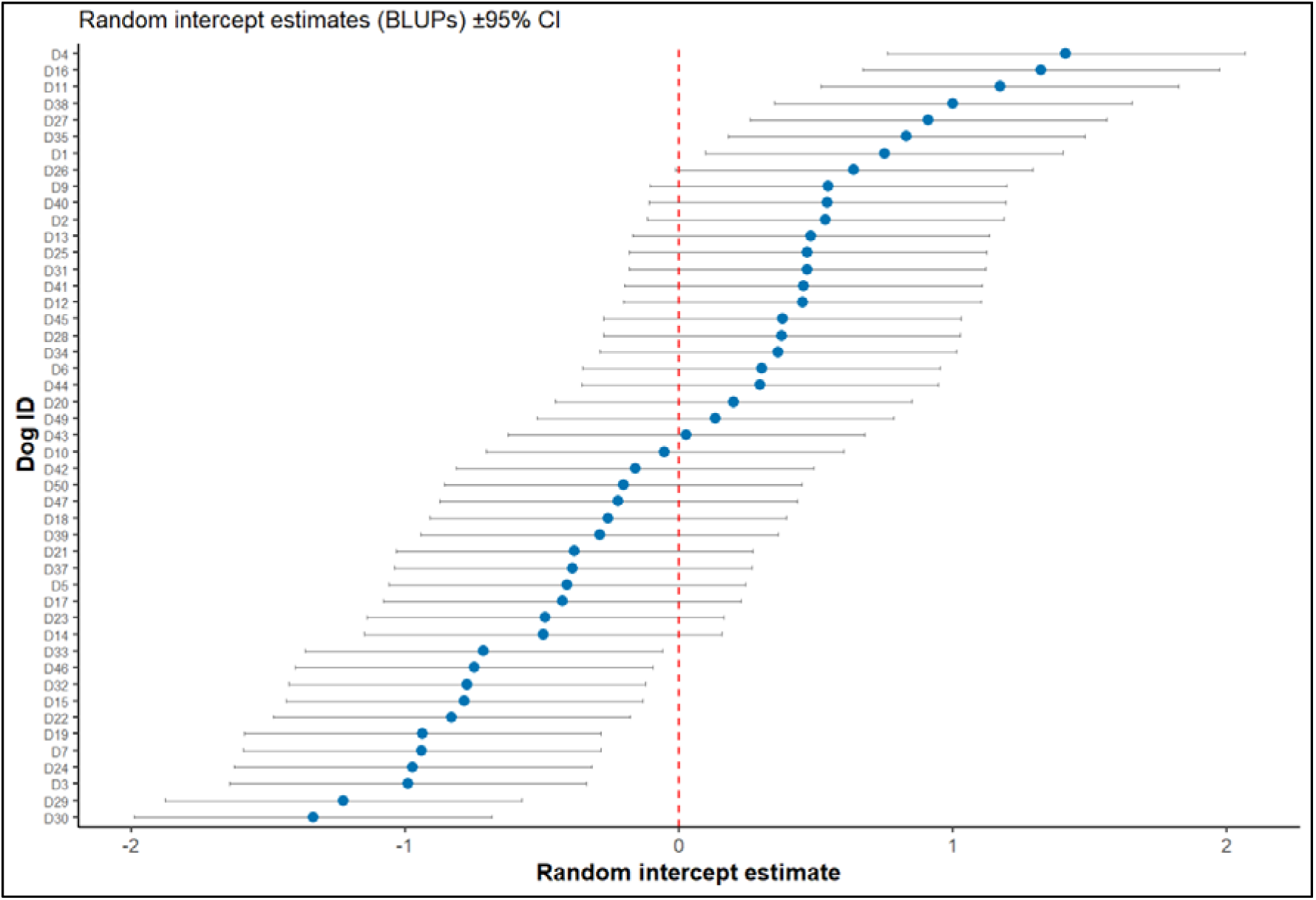
Random intercept estimates (BLUPs) for individual dogs from the linear mixed-effects model. Points represent individual-specific random intercepts, and horizontal bars indicate 95% confidence intervals.

### 2. Do free-ranging dogs modify their behavioural time investment across repeated trials?

To examine whether repeated exposure altered the allocation of time to different foraging behaviours, we fitted generalized linear mixed models (beta distribution) for each behavioural category, with proportional time investment as the response variable, Trial_number and Gender as fixed effects, and Dog_ID as a random intercept. For the behavioural time-investment analyses, the unit of analysis was the individual behavioural bout/event, not the dog-trial as a whole. Each event was assigned to a behavioural category, and its duration was expressed as a proportion of the total interaction time for that dog in that trial. Thus, the number of observations differed among behaviours and could exceed the number of dog-trial observations when a behaviour occurred multiple times within the same trial.

Across repeated exposures, we detected no significant trial-related change in the majority of strategic foraging behaviours. Time invested in sniffing (SN), multiple licking (ML), placing food on the ground (PG), licking (LI), rubbing (RB), carrying food (CF), foreleg use (FLU), dropping food on the ground (DG), and chewing (CW) did not vary significantly across trials or between sexes (all *p* > 0.05).

In contrast, repeated exposure selectively modified a subset of behaviours. Time spent eating increased significantly in Trial 3 (β = 0.55, *p* = 0.001) and remained elevated in Trial 4 (β = 0.36, *p* = 0.043) relative to Trial 1. Bowl-related manipulation showed some evidence of trial-related change. Nudging bowl behaviour increased in Trial 3, but this result should be interpreted cautiously because the behaviour was rare. UBNB showed a similar statistical pattern, but because it occurred in only six observations from five dogs, we treat this result as exploratory rather than as strong inferential evidence. Failed grab (FG) exhibited a transient shift, decreasing in Trial 3 (β = −0.69, *p* = 0.018) before increasing above baseline in Trial 4 (β = 0.49, *p* = 0.022). Similarly, head shaking (SH) increased significantly by Trial 4 (β = 0.19, *p* = 0.043).

Sex differences were evident for only a small subset of behaviours. Female dogs devoted more time to eating (ET) than males (β = −0.53, *p* = 0.013), whereas males spent more time performing upturning bowl by nudging (UBNB; β = 1.96, *p* = 0.001) and head shaking without food (SHNF; β = 1.00, *p* < 0.001). Apparent sex differences in rare behaviours such as UBNB and SHNF should be interpreted cautiously because these behaviours occurred in very few observations. Males also showed a marginal tendency to invest more time in failed grabs (FG; β = 0.57, *p* = 0.052), although this effect was not statistically significant.

Overall, these results indicate that repeated exposure did not broadly alter the dogs’ behavioural repertoire. Instead, behavioural adjustment was restricted to a small number of manipulative and consumption-related behaviours, while most measured behavioural components showed no detectable trial-related change. This pattern suggests selective refinement of key behavioural elements rather than wholesale changes in behavioural strategy (Supplementary information 2).

### 3. Do repeated exposures influence the time taken by free-ranging dogs to successfully consume food?

To determine whether repeated exposure influenced feeding efficiency (time from initial sniff to complete consumption), we fitted a mixed-effects Cox proportional hazards model with Trial_number and Gender as fixed effects and Dog_ID as a random intercept. Of the 188 trial observations, 134 ended in complete food consumption, while 54 trials were right-censored because the dog did not complete consumption within the 90-s observation period.

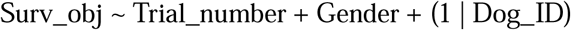

Repeated exposure significantly increased feeding efficiency. Relative to Trial 1, dogs were more likely to complete feeding sooner in Trial 2 (β = 1.03, hazard ratio [HR] = 2.79, z = 3.49, p < 0.001), with the strongest effect observed in Trial 3 (β = 1.53, HR = 4.60, z = 5.08, p < 0.001). Although the effect declined slightly in Trial 4, feeding completion remained significantly faster than in the initial exposure (β = 1.30, HR = 3.67, z = 4.44, p < 0.001) (Figure 3).

**Figure 3.**
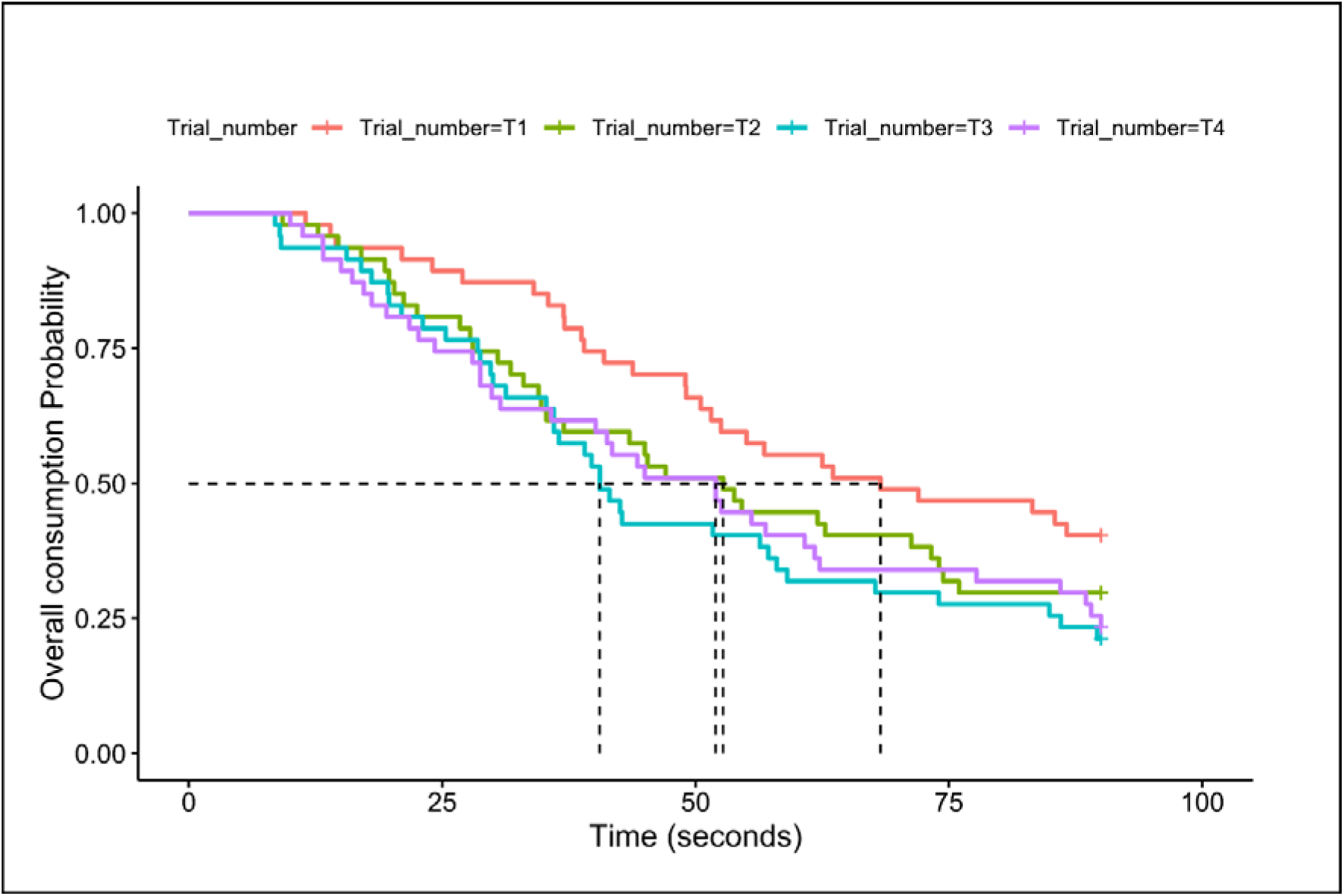
Feeding efficiency across experimental exposures. (a) Kaplan-Meier survival curves indicating the probability of overall food consumption over time (seconds) across the four trial phases (T1 to T4). Dashed drop-lines intersect the 50% consumption probability to illustrate the shift in median completion times across successive exposures (Supplementary information 3).

Sex also significantly influenced feeding efficiency. Male dogs had a lower hazard of completing feeding than females (β = −1.39, HR = 0.25, z = −2.14, p = 0.032), indicating that females completed the overall feeding process (from first sniff to final consumption) more rapidly than males. This is consistent with the earlier finding that females devoted proportionally more of their interaction time to eating itself rather than to preliminary sniffing or manipulation.

Overall, these results indicate that feeding efficiency improved after the first exposure: dogs completed consumption faster in Trials 2–4 than in Trial 1, with the strongest effect observed in Trial 3, while substantial among-dog variation in feeding completion remained.

### 4. What factors influence total interaction time?

To examine whether the latency to first interaction and location influenced the total interaction time, we fitted a Gamma generalized linear mixed model (GLMM) with a log link, using total interaction time as the response variable, Latency in seconds and Location as fixed effects, and Dog_ID as a random intercept (188 observations from 47 dogs).

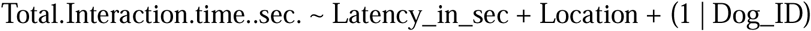

Longer latency to first interaction was associated with significantly longer total interaction times (β = 0.035, SE = 0.017, *z* = 2.11, *p* = 0.035) (Figure 4A). Relative to the reference location (Chandamari), dogs exhibited significantly shorter interaction times at Kalyani A Block (β = −0.880, SE = 0.437, *z* = −2.01, *p* = 0.044) and Rina Rd. (β = −1.838, SE = 0.586, *z* = −3.14, *p* = 0.002). Interaction times also tended to be shorter at Kalyani B Block (β = −0.846, SE = 0.432, *z* = −1.96, *p* = 0.050), Kalyani ITI More (β = −1.054, SE = 0.587, *z* = −1.79, *p* = 0.073), and Krishnapur (β = −0.977, SE = 0.592, *z* = −1.65, *p* = 0.099), whereas no significant differences were detected for the remaining locations (*p* > 0.10) (Figure 4B). Moderate among-individual variation was observed (Dog_ID variance = 0.067, SD = 0.258), with a residual dispersion of σ² = 0.417.

**Figure 4.**
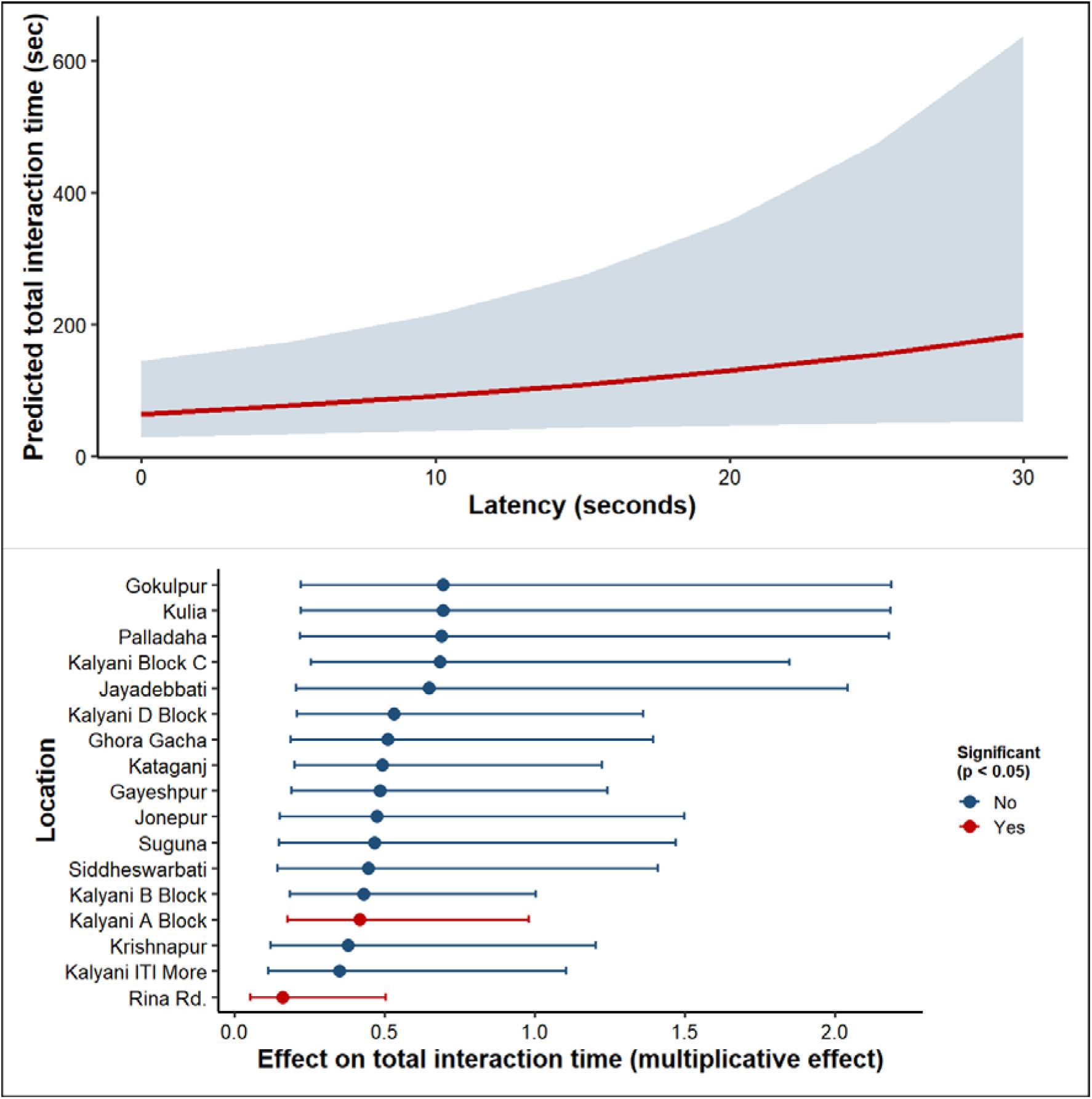
Predicted effects of latency and location on total interaction time from the Gamma generalized linear mixed-effects model (GLMM). **(A)** Population-level predicted relationship between latency and total interaction time, with the solid red line representing the fitted values and the shaded blue region indicating the 95% confidence interval. **(B)** Forest plot showing the relative effects of experimental locations on total interaction time compared with the reference location (Chandamari). Points represent exponentiated model coefficients (multiplicative effects), horizontal bars indicate 95% confidence intervals, and red points denote statistically significant effects (*p* < 0.05), whereas blue points indicate non-significant effects (Supplementary information 4).

Overall, these findings indicate that dogs taking longer to initiate interaction also spent longer completing the entire interaction sequence, while variation across locations suggests that spatial context contributed to variation in interaction time.

### 5. Do repeated exposures alter the decision-making time between olfactory inspection and licking behaviour?

To examine whether repeated exposure influenced the latency between the first sniff and first lick, we fitted a Gamma generalized linear mixed model (GLMM) with a log link, using sniff-to-lick latency as the response variable. Trial number and Location were included as fixed effects, with Dog_ID fitted as a random intercept (188 observations from 47 dogs).

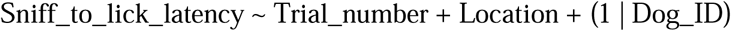

Repeated exposure significantly reduced sniff-to-lick latency. Relative to Trial 1, dogs transitioned from olfactory inspection to their first lick significantly faster in Trial 2 (β = −0.93, SE = 0.12, *z* = −7.74, *p* < 0.001), Trial 3 (β = −0.99, SE = 0.12, *z* = −8.30, *p* < 0.001), and Trial 4 (β = −0.85, SE = 0.12, *z* = −7.06, *p* < 0.001) (Figure 5A). Location also significantly influenced sniff-to-lick latency. Relative to the reference location, dogs at Gokulpur (β = 1.79, SE = 0.64, *z* = 2.80, *p* = 0.005) and Rina Rd. (β = 1.82, SE = 0.64, *z* = 2.85, *p* = 0.004) exhibited significantly longer sniff-to-lick latencies, whereas no significant differences were detected among the remaining study locations (*p* > 0.05) (Figure 5B). Moderate among-individual variation remained after accounting for the fixed effects (Dog_ID variance = 0.126, SD = 0.354), with a residual dispersion estimate of σ² = 0.311.

**Figure 5.**
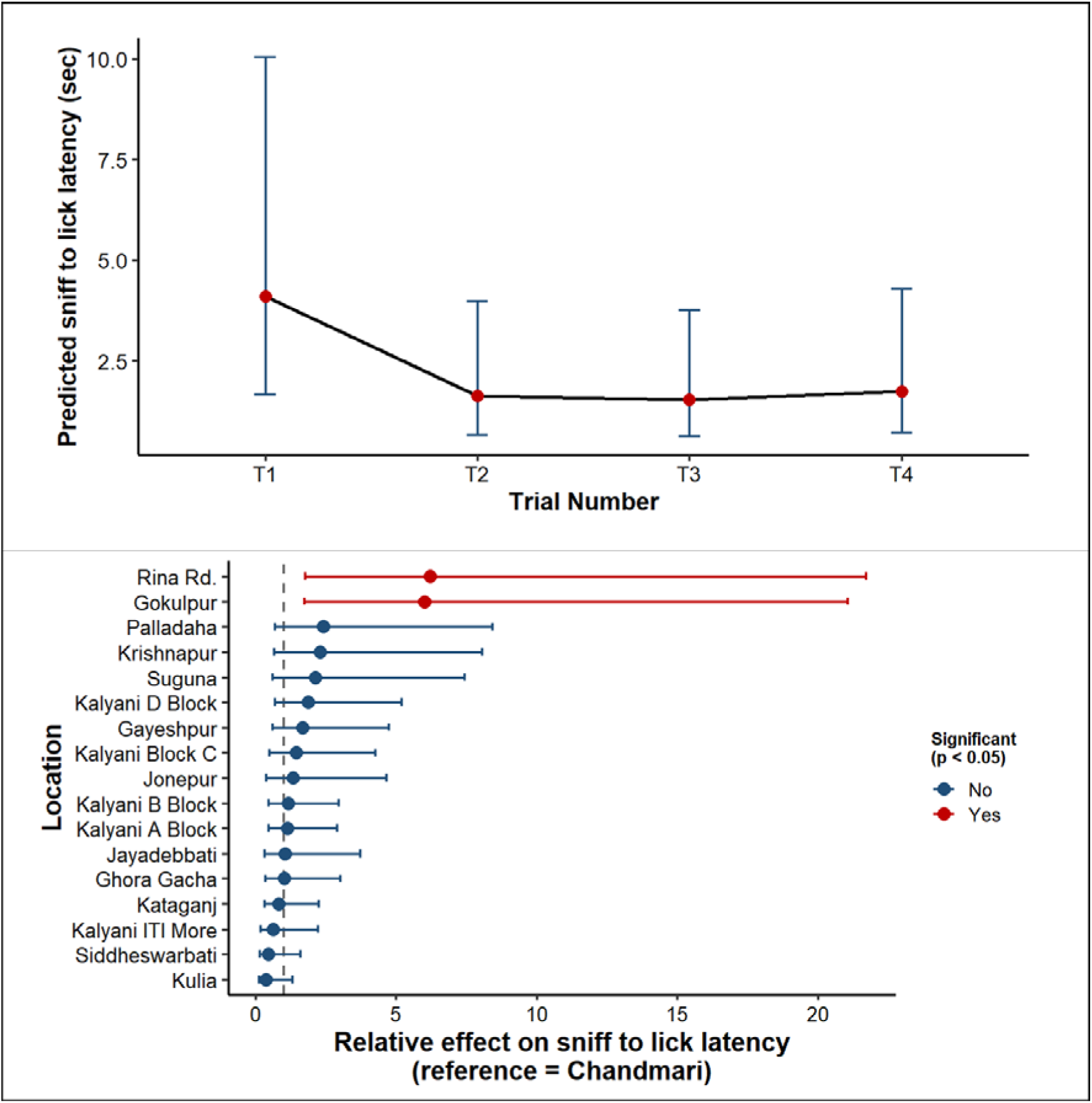
Predicted effects of trial number and location on the latency between the first sniff and first lick from the Gamma generalized linear mixed-effects model (GLMM). **(A)** Population-level predicted latency (±95% CI) across experimental trials (T1–T4). Points represent estimated marginal means and error bars denote 95% confidence intervals. **(B)** Forest plot showing the relative effects of experimental locations on sniff-to-lick latency compared with the reference location (Chandamari). Points represent exponentiated model coefficients (multiplicative effects), horizontal bars indicate 95% confidence intervals, and red points denote statistically significant effects (*p* < 0.05), whereas blue points indicate non-significant effects (Supplementary information 5).

Overall, these findings demonstrate that repeated exposure progressively shortened the interval between olfactory inspection and the first feeding attempt, indicating that dogs committed to feeding more rapidly with experience. In addition, differences among locations suggest that spatial context contributed to variation in post-inspection decision time.

### 6. Do individual free-ranging dogs show consistent behavioural strategies across repeated aversive foraging trials?

We conducted repeatability analysis, estimating repeatability coefficients (R) for each strategic behaviour. We used dog identity as a random effect, to assess whether behavioural variance was better explained by stable between-individual differences than by within-individual variation across trials (Nakagawa & Schielzeth, 2010). To capture consistency at two complementary levels, we estimated repeatability separately for whether a behaviour occurred at all (presence/absence, binomial-error rptR model, link-scale R) and for how much time was invested in it when it did occur (Gaussian linear mixed-effects models on log-transformed durations, log(x + 0.01)). In both cases, R was calculated as the proportion of total variance attributable to between-dog differences, with 95% bootstrap confidence intervals and permutation tests used to assess uncertainty and statistical support.

At the level of occurrence, several behaviours showed significant repeatability, indicating that individual dogs were consistent in whether they performed a given behaviour across trials. Eating, foreleg use, and head shaking showed the highest occurrence-based repeatability (R = 0.53–0.63, all *p* = 0.001), followed by carrying food, multiple licking, rubbing, chewing, sniffing, and placing food on ground, all of which were also significantly repeatable. Failed grab showed weaker but still significant repeatability. In contrast, licking, dropping food, nudging bowl, and head-shaking without food showed no significant occurrence-based repeatability (Table 1).

**Table 1.** Repeatability (R) of strategic behaviours across trials, estimated from duration-based and occurrence-based models. Duration-based estimates from Gaussian LMMs on log-durations; occurrence-based from binomial GLMMs on presence/absence data. Behaviours grouped by duration-based R: High (>0.40), Moderate (0.20–0.40), Low (<0.20).

| Sl No. | Behaviour | R (duration) | p (duration) | R (occurrence) | p (occurrence) |
| --- | --- | --- | --- | --- | --- |
| 1. | ET | 0.50 | <0.001 | 0.63 | 0.001 |
| 2. | FLU | 0.50 | <0.001 | 0.53 | 0.001 |
| 3. | SH | 0.48 | <0.001 | 0.53 | 0.001 |
| 4. | CF | 0.37 | <0.001 | 0.40 | 0.001 |
| 5. | ML | 0.34 | <0.001 | 0.37 | 0.001 |
| 6. | RB | 0.29 | <0.001 | 0.29 | 0.001 |
| 7. | CW | 0.27 | <0.001 | 0.24 | 0.001 |
| 8. | SN | 0.25 | <0.001 | 0.25 | 0.001 |
| 9. | PG | 0.23 | <0.001 | 0.24 | 0.001 |
| 10. | FG | 0.14 | 0.016 | 0.11 | 0.028 |
| 11. | LI | 0.07 | 0.106 | 0.07 | 0.075 |
| 12. | NB | 0.06 | 0.152 | 0.15 | 0.081 |
| 13. | DG | 0.05 | 0.190 | 0.06 | 0.153 |
| 14. | SHNF | 0.00 | 1.000 | 0.00 | 0.583 |

Note: Upturning bowl (UB) and upturn bowl by nudging (UBNB) occurred too rarely across trials to yield reliable repeatability estimates; both occurrence- and duration-based models either failed to converge or produced estimates driven by too few individuals to interpret. These two behaviours are therefore excluded from the table.

The same pattern emerged when consistency was instead assessed by the duration of behavioural investment. Eating time, foreleg use, and head shaking again showed the highest repeatability (R = 0.48–0.50, all *p* < 0.001), followed by carrying food, multiple licking, rubbing, chewing, sniffing, and placing food on ground, which were also significantly repeatable. Failed grab showed weaker but significant repeatability. Licking, nudging bowl, dropping food, and head shaking without food showed no significant duration-based repeatability (Table 1).

Together, these two independent measures converge almost completely on the same core set of behaviours. Eating, foreleg use, head shaking, carrying food, multiple licking, rubbing, sniffing, chewing, and placing food were consistently repeatable whether individual differences were assessed by whether a behaviour was performed or by how long it was performed, whereas licking, nudging bowl, dropping food, and head-shaking without food were consistently non-repeatable under both measures. This convergence indicates that the individual differences identified here reflect a genuine behavioural fingerprint rather than an artefact of a particular measurement scale. Free-ranging dogs therefore exhibit moderate but behaviour-specific consistency in their aversive foraging strategies. Behaviours central to resource handling and consumption (eating, foreleg use, head shaking) reflect stable individual tendencies, while behaviours such as licking, nudging, and dropping food show little evidence of stable individual differences, instead varying more within individuals or occurring too rarely to support reliable repeatability estimates (Figure 6).

**Figure 6.**
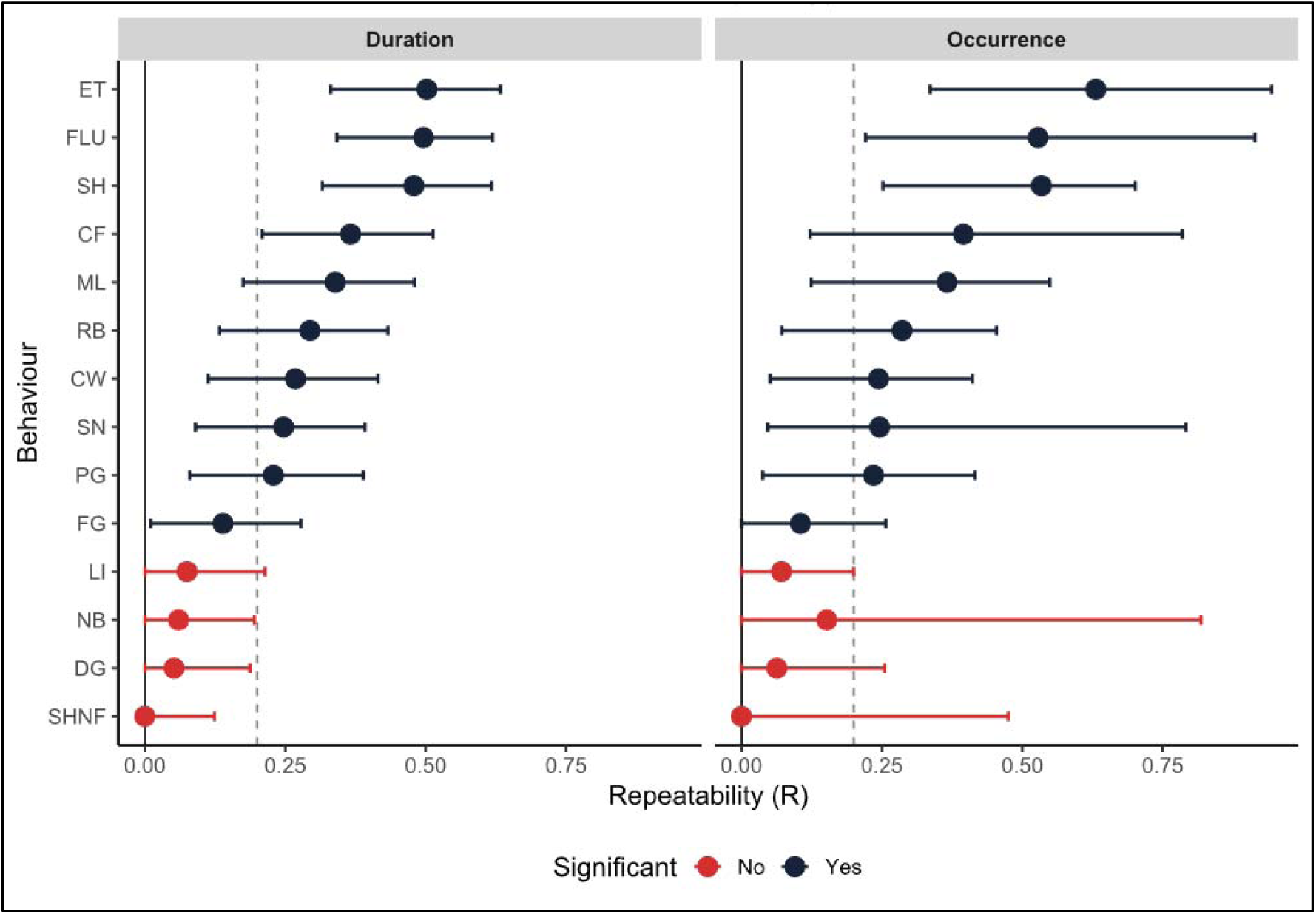
Behavioural repeatability across trials, assessed by duration and occurrence. Repeatability coefficients (R) with 95% bootstrap confidence intervals for each strategic behaviour, estimated separately from duration-based (left) and occurrence-based (right) repeatability models, with dog identity as a random effect. Dashed line marks R = 0.2 (threshold for low vs. moderate–high consistency). Dark points/bars indicate significant repeatability (*p* < 0.05); red indicates non-significant. Eating (ET), foreleg use (FLU), and head shaking (SH) show the highest, most consistent repeatability under both measures, while licking (LI), nudging bowl (NB), dropping food (DG), and head shaking without food (SHNF) show high within-individual flexibility under both.

### 7. Behavioural Sequence Consistency Across Repeated Trials

We quantified behavioural sequence consistency across trials (T1–T4) in 47 dogs using Optimal Matching (OM) distances with transition-rate-based substitution costs, yielding six within-individual trial-pair comparisons per dog (282 observations total).

Consecutive-trial distances declined over time: T1–T2 = 0.769 ± 0.283, T2–T3 = 0.726 ± 0.281, T3–T4 = 0.691 ± 0.298, indicating increasing sequence similarity in later trials. Non-consecutive comparisons showed the same pattern — distances involving Trial 1 remained high (T1–T3 = 0.809 ± 0.227; T1–T4 = 0.779 ± 0.236), while T2–T4 was lower (0.687 ± 0.281). A rank-based mixed-effects analysis confirmed a significant overall effect of trial-pair category (F = 4.438, *p* = 0.0007), with Tukey-adjusted post-hoc tests showing T1–T3 higher than both T2–T4 (*p* = 0.0025) and T3–T4 (*p* = 0.0066), and T1–T4 higher than T2–T4 (*p* = 0.0378). Because these six categories share trials, this is treated as descriptive evidence of temporal restructuring rather than a formal test of progressive change.

A within-dog directional permutation test on the consecutive comparisons (T1–T2, T2–T3, T3–T4) confirmed this trend directly, yielding a negative mean slope (−0.039 OM-distance units per step; *p* = 0.028); later trials were significantly more similar than earlier ones, though this does not establish strict monotonic improvement or identify the underlying mechanism (Figure 7).

**Figure 7.**
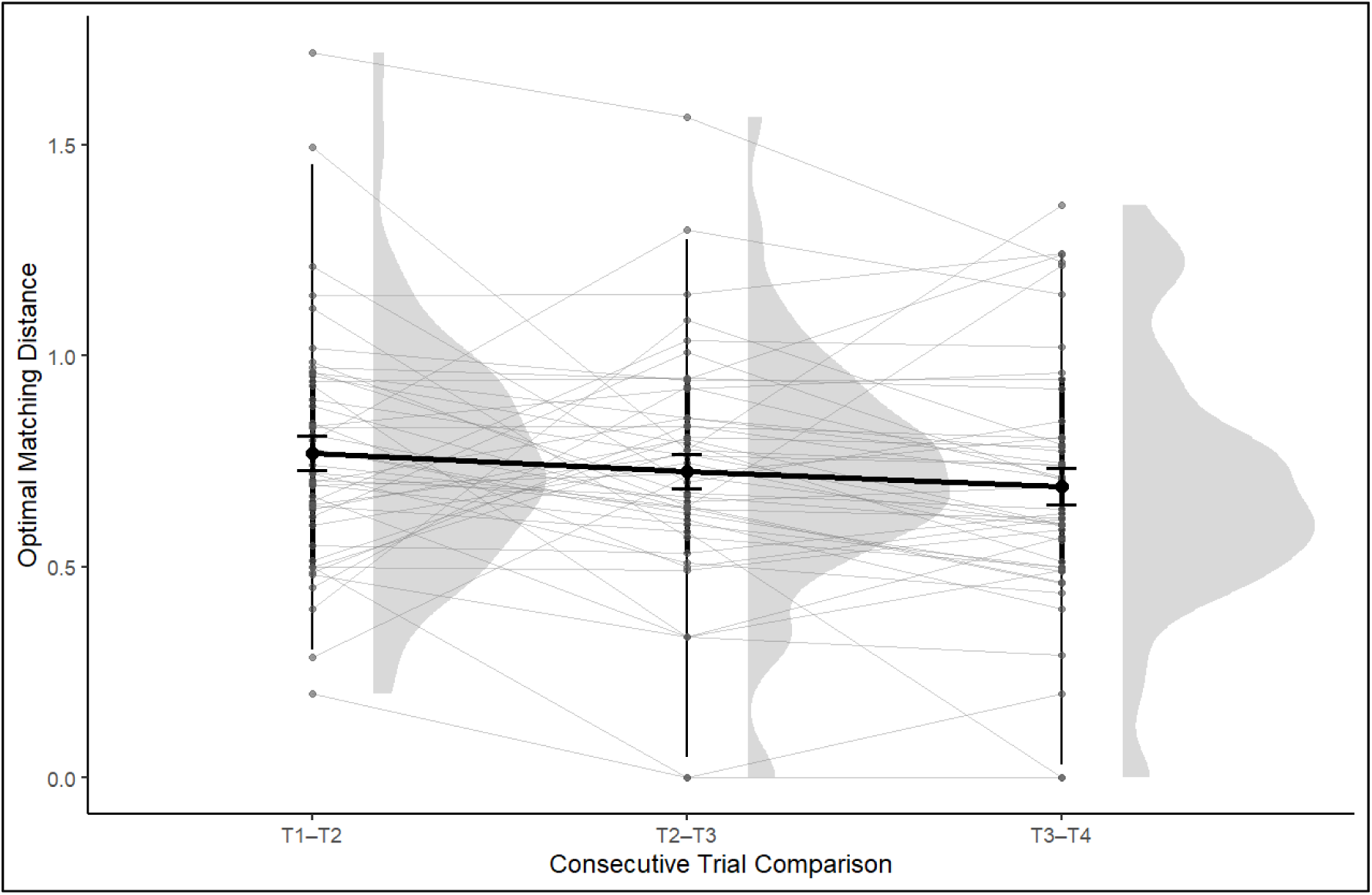
Raincloud trajectory plot of Optimal Matching (OM) distances across consecutive trial comparisons (T1–T2, T2–T3, and T3–T4). Half-violins depict the distribution of OM distances, points represent individual dogs, grey lines connect repeated observations within dogs, and the black line indicates the mean ± SE.

Substantial individual variation was also evident, with mean OM distances ranging from 0.133 (most consistent dog) to 1.231 (least consistent). Overall, dogs showed progressively more similar behavioural sequences across repeated exposures, alongside marked individual variation in the degree of this stabilisation (Supplementary information 6).

## Discussion

Free-ranging dogs did not overhaul their behavioural repertoire when they were repeatedly confronted with aversive stimuli along with edible food. Instead, they preserved core response tendencies while selectively refining the specific actions that determined how efficiently they resolved the challenge. By integrating behavioural frequencies, latency measures, interaction-time and location effects, duration-and occurrence-based repeatability analyses, survival-based feeding efficiency, and sequence structure, we found that repeated exposure reshaped particular decision points without eliminating individual behavioural tendencies (Bell et al., 2009; Nakagawa & Schielzeth, 2010). This balance between stability and selective adjustment points to a broader question in behavioural ecology (Dingemanse et al., 2010). The emerging question is how animals reconcile consistent individual dispositions with the flexibility needed to cope with recurring, ecologically relevant challenges in human-dominated environments. This supports our third hypothesis: partially independent mechanisms for risk assessment and handling, rather than uniform stability or broad flexibility.

The first key pattern concerned the timing of adjustment. Approach latency did not decline across trials, nor did it differ between sexes; instead, it was highly repeatable across individuals, with roughly half of the total variance attributable to consistent differences among dogs rather than to trial number. This is a distinct repeatability estimate from the behaviour-specific values reported for strategic foraging behaviours such as eating, foreleg use, and head shaking, which were independently repeatable whether individual differences were measured by duration of engagement or by simple occurrence across trials. That convergence across two independent measurement scales indicates that the consistency in strategic foraging behaviours reflects genuine trait-like individual dispositions rather than an artefact of how behavioural investment happened to be quantified. Taken together with the approach-latency result, this suggests that individual consistency in this system is not confined to a single behavioural measure but is detectable across both the decision to engage with the resource and the specific strategies used once engagement occurred. Allowing individual dogs to vary in how their latency to approach changed across trials did not improve model fit, indicating that dogs did not differ from one another in the direction or extent of any trial-related adjustment. This shows that the decision to approach the resource was shaped more by stable individual disposition than by prior experience within the experiment. Biologically, this is meaningful because it separates the initial decision to approach a risky resource from the subsequent decisions involved in evaluating and consuming it: approach behaviour appears to function as a consistent trait-like response, largely fixed at the level of the individual, whereas the timing and efficiency of downstream feeding decisions changed measurably with experience. Any learning-like adjustment therefore occurred mainly downstream of first contact, rather than in the approach phase itself. This distinction aligns with frameworks that view exploration–avoidance tendencies and proactive–reactive coping styles as separable behavioural axes (Réale et al., 2007; Sih et al., 2004). It also reflects a broader pattern across taxa, in which risk assessment and subsequent resource handling are governed by at least partially independent mechanisms (Sih & Bell, 2008). Rather than being specific to dogs or this foraging paradigm, the results point to a general principle: repeated exposure to a challenge need not affect the full behavioural sequence uniformly, but can instead leave early, dispositional decision points stable while selectively reshaping the decision nodes that follow.

The second key pattern was that this selective adjustment translated into functional gains. Feeding efficiency improved sharply from Trial 1, peaking at Trial 3, with a slight easing by Trial 4 that nonetheless remained far above baseline; the transition from sniffing the food to licking it followed the same pattern, dropping sharply after the first exposure, reaching its shortest latency at Trial 3, and remaining substantially reduced at Trial 4. Certain eating-related actions, such as eating behaviour itself and nudging the bowl, also changed selectively across trials. Total interaction time was not modelled as a function of trial number, but was instead predicted by how long a dog took to first engage with the setup and by the specific location of testing: dogs that approached more cautiously subsequently spent longer interacting with the food, and interaction times varied across sites independently of experience. Therefore, dogs became more effective and efficient at finishing the task without turning uniformly less cautious from the start or overhauling their entire behavioural approach; instead, individual differences in initial caution carried through into how long the interaction itself lasted.

In anthropogenic environments, aversive food, spiced, decaying, or embedded in refuse, is often abundant and predictable, whereas alternative food sources may be scarcer or less reliable (Garvey et al., 2020). Why did dogs refine their handling of an aversive resource rather than abandoning it, given that their underlying willingness to approach it remained stable regardless of experience? Under these conditions, persisting with the resource while minimizing the cost of interacting with it may represent an optimal foraging solution: refinement of handling reduces the perceived cost of aversion relative to the caloric benefit, rather than requiring dogs to forgo the resource altogether. Looked at this way, the dogs’ selective adjustments represent an adaptive strategy for maximizing energetic gain despite the persistent cost imposed by the aversive stimulus (Mathot et al., 2012), consistent with the optimal-foraging logic introduced earlier: dogs tolerated the aversive cost because the caloric payoff justified it (Sih & Christensen, 2001; Sih & Bell, 2008). Given that such food recurs reliably in the scavenging environment of free-ranging dogs, this refinement strategy would be adaptive over repeated encounters.

The findings on interaction duration and the sniff-to-lick transition together clarify where, within the foraging sequence, dogs’ experience-dependent adjustments were concentrated. Sniff-to-lick latency shortened sharply with repeated exposure, whereas total interaction time was not itself predicted by trial number, but instead varied with each individual’s initial approach latency and with the testing location. Taken together, these results suggest that repeated exposure sped up the specific transition from inspection to feeding commitment, without producing a parallel, trial-driven change in how long dogs spent engaging with the setup overall. Instead, the duration of that overall engagement appears tied to a more stable individual trait: dogs that approached more cautiously also tended to spend longer manipulating and evaluating the food once they arrived, suggesting that initial caution and prolonged handling represent linked aspects of an individual’s response to the challenge, distinct from the specific, trial-sensitive speeding-up of the sniff-to-lick decision itself.

To our knowledge, this is among the first studies in free-ranging dogs to jointly examine behavioural consistency, temporal efficiency, and sequence structure within a repeated-exposure paradigm, providing a more integrated understanding of behavioural organisation than frequency-based measures alone (Garland et al., 2012). However, the study was limited to four exposures, precluding assessment of long-term behavioural stability. Although approach latency and strategic foraging behaviours both showed high repeatability, it remains unknown whether these consistencies persist across longer timescales or different contexts. The environmental factors underlying location effects and the relative contributions of associative learning versus non-associative habituation could not be resolved. Notably, site-level flux did not significantly predict either measure, ruling out human/vehicle activity as the explanation. The observed differences could be a result of specific exposures to certain food items in the individual localities, or the chance testing of individuals of a certain personality type at a higher frequency, factors that could not be controlled for in this experimental set-up.

Rare manipulative behaviours (e.g., UB, UBNB) limited inference for some behavioural categories. Future studies should extend repeated testing over longer periods, experimentally investigate the environmental drivers of spatial effects, and integrate sequence-based analyses with physiological or cognitive measures to better distinguish learning processes from habituation.

Taken together, these findings indicate that free-ranging dogs respond to repeated aversive foraging exposure through targeted behavioural refinement rather than wholesale change, and that this refinement operates within, not against, stable individual response patterns. More broadly, our findings suggest that behavioural adaptation need not involve reorganization of an entire behavioural repertoire (Mazza & Šlipogor, 2024). Instead, repeated experience may often modify a limited number of critical decision points while leaving the broader behavioural architecture intact. This offers a case study in how animals reconcile behavioural consistency with adaptive flexibility and, more specifically, how they weigh persistent costs against reliable caloric returns, when navigating recurring, ecologically realistic challenges in the anthropogenic environments they increasingly inhabit.

## Ethical statement

The study design did not violate the Animal Ethics regulations of the Government of India (Prevention of Cruelty to Animals Act 1960, Amendment 1982). The protocol for the experiment was approved by the IISER Kolkata Animal Ethics Committee.

## Authors’ contributions

TSP and AB conceptualised and designed the study. TSP along with SB conducted fieldwork. TSP and SB were involved in data decoding, data cleaning and data analysis. TSP wrote the manuscript; SB helped in writing and experimental diagram making. AB supervised the project, offered continuous input, and critically reviewed the manuscript.

## Acknowledgement

We thank all members of the BEL Lab for their valuable insights and support throughout the study. We are especially grateful to Imran Mondal for his assistance during fieldwork.

## Data availability

Will be made available on publication of the manuscript after peer review.

## Ethical approval

The study design complied with the Animal Ethics regulations of the Government of India (Prevention of Cruelty to Animals Act, 1960, as amended in 1982). The experimental protocol was approved by the IISER Kolkata Animal Ethics Committee as part of a larger project funded by SERB (EMR/2016/000595).

## References

1. Akaike, H. (1974). A new look at the statistical model identification. IEEE Transactions on Automatic Control, 19(6), 716–723. IEEE Xplore Entry.

2. Balleine, B. W., & Dickinson, A. (1998). Goal-directed instrumental action: contingency and incentive learning and their cortical substrates. Neuropharmacology, 37(4-5), 407–419.

3. Bell, A. M., Hankison, S. J., & Laskowski, K. L. (2009). The repeatability of behaviour: a meta-analysis. Animal behaviour, 77(4), 771–783.

4. Bhattacharjee, D., & Bhadra, A. (2020). Humans dominate the social interaction networks of urban free-ranging dogs in India. Frontiers in Psychology, 11, 2153.

5. Bhattacharjee, D., Sarkar, R., Sau, S., & Bhadra, A. (2021). Sociability of Indian free-ranging dogs (Canis lupus familiaris) varies with human movement in urban areas. Journal of comparative psychology, 135(1), 89.

6. Bhattacharjee, D., Sau, S., Das, J., & Bhadra, A. (2017). Free-ranging dogs prefer petting over food in repeated interactions with unfamiliar humans. Journal of Experimental Biology, 220(24), 4654–4660.

7. Biswas, S., Nandi, S., Pal, T. S., Lahiri, A., Roy, A., Gope H, Ghosh K, Touhid S, Ghosh S, Bhadra A. (2026). Begging for morsels: social context and sex shape begging in free-ranging dogs. Animal Behaviour, 240, 123693. 10.1016/j.anbehav.2026.123693

8. Bradshaw, J. W. (2006). The evolutionary basis for the feeding behavior of domestic dogs (Canis familiaris) and cats (Felis catus). The Journal of nutrition, 136(7), 1927S–1931S.

9. Budaev, S., & Brown, C. (2011). Personality traits and behaviour. Fish cognition and behavior, 135–165.

10. Burritt, E. A., & Provenza, F. D. (1991). Ability of lambs to learn with a delay between food consumption and consequences. Applied Animal Behaviour Science, 32(2-3), 221– 232.

11. Butler, J. R., Brown, W. Y., & Du Toit, J. T. (2018). Anthropogenic food subsidy to a commensal carnivore: the value and supply of human faeces in the diet of free-ranging dogs. Animals, 8(5), 67.

12. Coleman, K., & Wilson, D. S. (1998). Shyness and boldness in pumpkinseed sunfish: individual differences are context-specific. Animal behaviour, 56(4), 927–936.

13. Dias, R., Robbins, T. W., & Roberts, A. C. (1996). Dissociation in prefrontal cortex of affective and attentional shifts. Nature, 380(6569), 69–72.

14. Dingemanse, N. J., Kazem, A. J., Réale, D., & Wright, J. (2010). Behavioural reaction norms: animal personality meets individual plasticity. Trends in ecology & evolution, 25(2), 81–89.

15. Fragoso, F. P., & Brunet, J. (2023). Honey bees exhibit greater patch fidelity than bumble bees when foraging in a common environment. Ecosphere, 14(6), e4606.

16. Frank, H. E., Amato, K., Trautwein, M., Maia, P., Liman, E. R., Nichols, L. M., & Dunn, R. R. (2022). The evolution of sour taste. Proceedings of the Royal Society B, 289(1968), 20211918.

17. Gabadinho, A., Ritschard, G., Müller, N. S., & Studer, M. (2011). Analyzing and visualizing state sequences in R with TraMineR. Journal of statistical software, 40, 1–37.

18. Garland, E. C., Lilley, M. S., Goldizen, A. W., Rekdahl, M. L., Garrigue, C., & Noad, M. J. (2012). Improved versions of the Levenshtein distance method for comparing sequence information in animals’ vocalisations: tests using humpback whale song. Behaviour, 149(13-14), 1413–1441.

19. Garvey, P. M., Banks, P. B., Suraci, J. P., Bodey, T. W., Glen, A. S., Jones, C. J., … & Sih, A. (2020). Leveraging motivations, personality, and sensory cues for vertebrate pest management. Trends in ecology & evolution, 35(11), 990–1000.

20. Gosling, S. D. (2001). From mice to men: what can we learn about personality from animal research?. Psychological bulletin, 127(1), 45.

21. Hughes, R. N., Kaiser, M. J., Mackney, P. A., & Warburton, K. (1992). Optimizing foraging behaviour through learning. Journal of Fish Biology, 41, 77–91.

22. J. Valone, T. (2006). Are animals capable of Bayesian updating? An empirical review. Oikos, 112(2), 252–259.

23. Lyons, D. M., Price, E. O., & Moberg, G. P. (1988). Individual differences in temperament of domestic dairy goats: constancy and change. Animal Behaviour, 36(5), 1323–1333.

24. Mathot, K. J., Wright, J., Kempenaers, B., & Dingemanse, N. J. (2012). Adaptive strategies for managing uncertainty may explain personality-related differences in behavioural plasticity. Oikos, 121(2), 210–220.

25. Mazza, V., & Šlipogor, V. (2024). Behavioral flexibility and novel environments: integrating current perspectives for future directions. Current zoology, 70(3), 304–309.

26. Mehiel, R., & Bolles, R. C. (1988). Learned flavor preferences based on calories are independent of initial hedonic value. Animal Learning & Behavior, 16(4), 383–387.

27. Nakagawa, S., & Schielzeth, H. (2010). Repeatability for Gaussian and non Gaussian data: a practical guide for biologists. Biological Reviews, 85(4), 935–956.

28. Nandi, S., Lahiri, A., Pal, T. S., Roy, A., Bairagya, R., & Bhadra, A. (2026). Treats or affection? Understanding reward preferences in Indian free-ranging dogs. Animal Cognition, 29(1), 31. 10.1007/s10071-026-02046-4

29. Pal, T. S. (2022). Understanding the Preference of Citrus Food among Free-ranging Dogs (Doctoral dissertation, Indian Institute of Science Education and Research Kolkata).

30. Pal, T. S., Debnath, P., Biswas, S., & Bhadra, A. (2026). Lemons and learning: Juvenile free ranging dogs are more persistent but less selective foragers than adults. Journal of Zoology.

31. Pal, T. S., Nandi, S., Gope, H., Malakar, A., Sarkar, R., Biswas, S., & Bhadra, A. (2025b). Exploring Scavenging Strategies and Cognitive Problem-Solving in Indian Free-Ranging Dogs. *arXiv preprint arXiv:2512.01637*.

32. Pal, T. S., Nandi, S., Sarkar, R., & Bhadra, A. (2025a). When Life Gives You Lemons, Squeeze Your Way Through: Understanding Citrus Avoidance Behaviour by Free-Ranging Dogs in India. Applied Animal Behaviour Science, 106682.

33. Payne, R. J., & Pagel, M. (1997). Why do animals repeat displays?. Animal behaviour, 54(1), 109–119.

34. Provenza, F. D. (1996). Acquired aversions as the basis for varied diets of ruminants foraging on rangelands. Journal of animal science, 74(8), 2010–2020.

35. R Core Team. 2022. R: A language and environment for statistical computing. R Foundation for Statistical Computing, Vienna, Austria. https://www.R-project.org/

36. Réale, D., Reader, S. M., Sol, D., McDougall, P. T., & Dingemanse, N. J. (2007). Integrating animal temperament within ecology and evolution. Biological reviews, 82(2), 291–318.

37. Sarkar, R., & Bhadra, A. (2022). How do animals navigate the urban jungle? A review of cognition in urban-adapted animals. Current Opinion in Behavioral Sciences, 46, 101177.

38. Sarkar, R., Bhowmick, A., Dasgupta, D., Banerjee, R., Chakraborty, P., Nayek, A., … & Bhadra, A. (2023). Eating smart: Free-ranging dogs follow an optimal foraging strategy while scavenging in groups. Frontiers in Ecology and Evolution, 11, 1099543.

39. Sarkar, R., Maji, S., Pal, T. S., Rajratna, A. D., Ghosh, A., Roy, M., … & Bhadra, A. (2025a). Trick or Treat? Free-ranging dogs use human behavioural cues for foraging. arXiv preprint arXiv:2512.18681.

40. Sarkar, R., Pal, T. S., Maji, S., Nandi, S., Lahiri, A., Basumatary, A. K., … & Bhadra, A. (2025b). Sight, smell and more: What cues do free-ranging dogs use for decision-making while scavenging?. arXiv preprint arXiv:2512.00058.

41. Sarkar, R., Pal, T. S., Murmu, S., & Bhadra, A. (2024). The mouth speaks as much as the eyes: Free-ranging dogs depend on inner facial features for human recognition. 10.48550/arXiv.2407.07192.

42. Sarkar, R., Sau, S., & Bhadra, A. (2019). Scavengers can be choosers: A study on food preference in free-ranging dogs. Applied Animal Behaviour Science, 216, 38–44.

43. Schwarz, G. (1978). Estimating the dimension of a model. The Annals of Statistics, 6(2), 461–464. Project Euclid Entry.

44. Sclafani, A. (1995). How food preferences are learned: Mechanisms and methods. Science, 267(5203), 1548–1549.

45. Sih, A., & Bell, A. M. (2008). Insights for behavioral ecology from behavioral syndromes. Advances in the Study of Behavior, 38, 227–281.

46. Sih, A., & Christensen, B. (2001). Optimal diet theory: when does it work, and when and why does it fail?. Animal behaviour, 61(2), 379–390.

47. Sih, A., & Del Giudice, M. (2012). Linking behavioural syndromes and cognition: a behavioural ecology perspective. Philosophical Transactions of the Royal Society B: Biological Sciences, 367(1603), 2762–2772.

48. Sih, A., Bell, A., & Johnson, J. C. (2004). Behavioral syndromes: an ecological and evolutionary overview. Trends in ecology & evolution, 19(7), 372–378.

49. Stagner, J. P., Laude, J. R., & Zentall, T. R. (2012). Pigeons prefer discriminative stimuli independently of the overall probability of reinforcement and of the number of presentations of the conditioned reinforcer. Journal of Experimental Psychology: Animal Behavior Processes, 38(4), 446.

50. Stephens, D. W., Brown, J. S., & Ydenberg, R. C. (Eds.). (2008). Foraging: behavior and ecology. University of Chicago Press.

51. Stoffel, M. A., Nakagawa, S., & Schielzeth, H. (2017). rptR: repeatability estimation and variance decomposition by generalized linear mixed effects models. Methods in ecology and evolution, 8(11), 1639–1644.

52. Sugrue, L. P., Corrado, G. S., & Newsome, W. T. (2004). Matching behavior and the representation of value in the parietal cortex. science, 304(5678), 1782–1787.

53. Svartberg, K., Tapper, I., Temrin, H., Radesäter, T., & Thorman, S. (2005). Consistency of personality traits in dogs. Animal Behaviour, 69(2), 283–291.

54. Tolman, E. C., Ritchie, B. F., & Kalish, D. (1946). Studies in spatial learning. II. Place learning versus response learning. Journal of experimental psychology, 36(3), 221.

55. Woodford, R. (2012). Feed your best friend better: Easy, nutritious meals and treats for dogs. Andrews McMeel Publishing.

56. Zuckerman, M. (1991). Psychobiology of personality (Vol. 10). Cambridge University Press.

