## Supplemental file for "Fixed Players, Fluid Moves: Refining Strategy While Preserving Identity in Free-Ranging Dogs"

Supplementary figure 1.

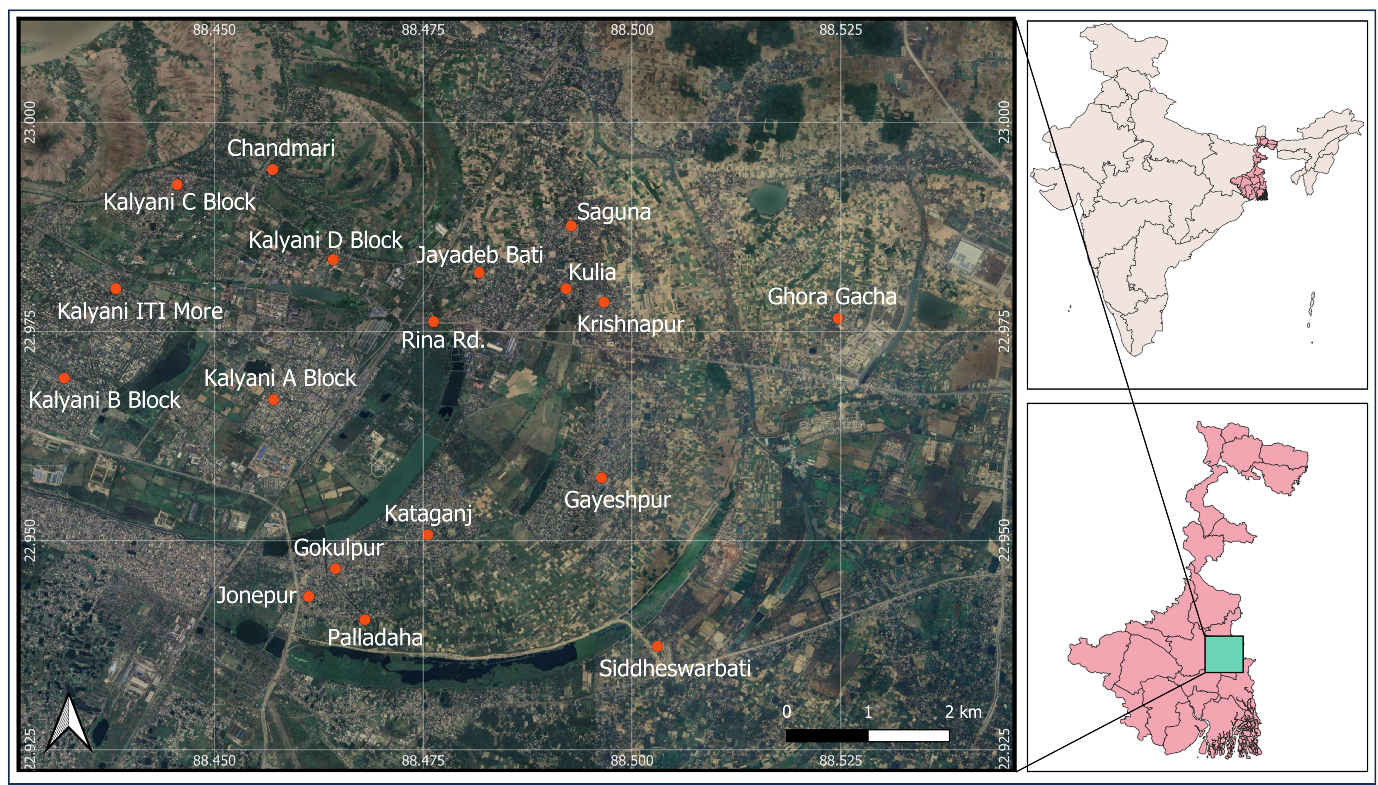

Fig. A map showing the sampling sites for the experiments in the Nadia district of West Bengal, India.

**Supplementary table 1:**

| **Name of the Behaviour** | **Behavioural**  **Code** | **Definition** |
| --- | --- | --- |
| Upturning bowl by using teeth | UB | The act of grasping a bowl with teeth and partially or fully flipping it over. The act of carrying the bowl of food and some spillage of liquid or food or both. |
| Head shake | SH | The rapid sideways movement of the head following the act of grabbing food in the mouth. |
| Head shake without food | SHNF | The rapid sideways movement of the head without food in the mouth. |
| Placing food on the ground | PG | The act of placing food on the ground in a courteous manner, ensuring it's not dropped but set down gently. |
| Drop food on the ground | DG | Dropping food from a height onto the ground, not employing a gentle movement. |
| Rubbing food on the ground | RB | Rub the food against the ground using teeth, mouth, or snout. Dogs can drag the chicken piece using teeth/mouth on the ground. |
| Using foreleg | FLU (FLL, FLR, FLB) | The action of manipulating food using one or more forelegs (left, right, or both) to free food or tamper the food or tamper the bowl. |
| Multiple lick | ML | The action of repetitively licking using the tongue, occurring more than once, typically two or more times. |
| Nudging bowl | NB | A behaviour where dog nudges the bowl containing food, using its snout, typically resulting in a mild movement or displacement of the bowl. |
| Upturn bowl by nudging | UBNB | The act of overturning a bowl by applying force through a gentle nudge, resulting in a complete or partial reversal of its position rather than just a mild displacement. |
| Only Chewing | CW | The action of a dog using its mouth to bite, gnaw, or chew on various objects such as toys, bones, or food items. Chewing but after that not eating the food. If chewing and then eat the food then it will not be considered as chewing. |
| Eat | ET | The act of consuming food entirely, including consuming all parts together. If chewing and then after chewing eating the whole food, it will be counted as eating. |
| Partial eating | PE | Specific eating behaviour when instead of eating entire chicken piece, parts of the chicken piece is being eaten one at a time. If a chicken piece is being chewing and eating, it will be considered as PE. |
| Sniff | SN | The dog approaches the bowl and investigates it by sniffing from a distance of within 10cm. |
| Lick | LI | The act of a dog using its tongue to clean or consume food from a bowl. This behaviour excludes the use of the muzzle or snout. |
| Carrying food or bowl of food or bowl | CF | The act of picking of the chicken piece from the bowl and walking/ trotting/running away from the bowl or from the ground with chicken piece inside its mouth. It can be positional displacement of the food from the bowl. The act of carrying the bowl of food, bowl and place it on the ground. The act of carrying food. |
| Failed grab | FG | This category includes all the situations in which the dog tries to take up the chicken with its mouth but cannot manage to do so. This may entail making tries in the bowl or on the ground. It eventually drops the food several times on the ground or bowl. |
| Not visible | NV | When dog’s activity is not detectable and visible from the decoder’s point of view. |

Supplementary information:

Supplementary information 1:

**Model Summary**

| Metric | Value |
| --- | --- |
| **Model** | Gaussian (identity link) |
| **Formula** | log_Latency ~ Trial_number + Gender + (1 \| Dog_ID) |
| **Data** | data880 |
| **AIC** | 507.1 |
| **BIC** | 529.8 |
| **logLik** | -246.6 |
| **Deviance** | 493.1 |
| **df.resid** | 181 |
| **Dispersion (σ²)** | 0.529 |

**Random Effects**

| Group | Variable | Variance | [Std.Dev](https://std.dev/). |
| --- | --- | --- | --- |
| Dog_ID | Intercept | 0.5830 | 0.7635 |
| Residual | - | 0.5289 | 0.7273 |

**Number of obs:** 188, **groups:** Dog_ID (47)

**Fixed Effects**

| Predictor | Estimate | Std. Error | z value | Pr(>\|z\|) | Signif. |
| --- | --- | --- | --- | --- | --- |
| (Intercept) | 0.3510 | 0.1797 | 1.953 | 0.0508 | . |
| Trial_numberT2 | 0.1450 | 0.1500 | 0.967 | 0.3337 |  |
| Trial_numberT3 | -0.1616 | 0.1500 | -1.077 | 0.2815 |  |
| Trial_numberT4 | -0.0038 | 0.1500 | -0.025 | 0.9801 |  |
| GenderM | -0.0822 | 0.2567 | -0.320 | 0.7489 |  |

**Significance codes:** *** p < 0.001, ** p < 0.01, * p < 0.05, . p < 0.1

**Repeatability of approach latency**

- Repeatability (R) = **0.531** ± 0.074
- 95% Confidence Interval (CI) = **0.362–0.651**
- Likelihood-ratio test (LRT): **P = 3.99 × 10⁻¹⁵**
- Permutation test: **P = 0.001**

Approach latency exhibited significant repeatability across repeated trials, with approximately 53.1% of the total variance attributable to consistent differences among individual dogs.

Supplementary information 2:

| **Strategic Behaviour** | **N (Obs/Groups)** | **Dispersion** | **Intercept** | **T2** | **T3** | **T4** | **GenderM** | **Random Var** |
| --- | --- | --- | --- | --- | --- | --- | --- | --- |
| ML | 233/42 | 6.95 | -1.6148*** | 0.0523 | 0.0567 | 0.0115 | -0.0787 | 0.5306 |
| ET | 133/41 | 9.09 | -0.8171*** | 0.1471 | 0.5527** | 0.3571* | -0.5263* | 0.2401 |
| FG | 88/34 | 31.0 | -3.1511*** | -0.0260 | -0.6944* | 0.4899* | 0.5725 | 0.4579 |
| SH | 326/40 | 85.7 | -3.4299*** | 0.0989 | 0.0315 | 0.1906* | -0.1392 | 0.2189 |
| PG | 201/45 | 58.3 | -3.5265*** | 0.2038 | 0.1382 | 0.0296 | -0.0696 | 0.1168 |
| LI | 120/43 | 7.34 | -2.6642*** | 0.0714 | 0.0045 | 0.2101 | 0.1326 | ~0 |
| RB | 221/40 | 11.2 | -1.7725*** | -0.0680 | -0.0382 | -0.0721 | -0.0889 | 0.0292 |
| CF | 237/45 | 18.1 | -2.5945*** | 0.1121 | 0.1274 | 0.1876 | -0.1520 | 0.0871 |
| CW | 86/33 | 11.7 | -1.8547*** | 0.4403 | -0.0988 | 0.1331 | 0.2629 | 0.0980 |
| FLU | 94/22 | 17.1 | -2.6208*** | -0.1226 | 0.1618 | -0.1958 | 0.0194 | 0.0095 |
| DG | 28/18 | 39.4 | -3.8407*** | 0.5073 | 0.2624 | -0.2967 | 0.0082 | ~0 |
| NB | 10/8 | 41.5 | -3.1284*** | -1.0877 | 1.1844** | -0.1668 | 0.4616 | ~0 |
| SHNF | 9/7 | 239.0 | -3.7665*** | -0.3939 | -0.1593 | - | 0.9960*** | ~0 |
| UBNB | 6/5 | 50.3 | -4.3273*** | 0.1352 | 1.7307*** | - | 1.9577** | ~0 |
| SN | 33/16 | 1.97 | -0.9747* | -0.4810 | 0.0229 | -0.2476 | -0.0722 | ~0 |

- Significance codes: ***p<0.001, **p<0.01, *p<0.05
- T2, T3, T4 = Trial numbers compared to reference (T1)
- ~0 indicates variance near zero (< 0.001)
- Dash (-) indicates no data for that trial
- Models sorted by sample size (descending) for easier comparison

Supplementary information 3:

**Model Summary**

| Metric | Value |
| --- | --- |
| **Model** | Mixed effects coxme model |
| **Formula** | surv_obj ~ Trial_number + Gender + (1 \| Dog_ID) |
| **Data** | data |
| **Events, n** | 134, 188 |

**Random Effects**

| Group | Variable | SD | Variance |
| --- | --- | --- | --- |
| Dog_ID | Intercept | 1.992 | 3.968 |

**Model Fit Statistics**

| Statistic | Value | df | p | AIC | BIC |
| --- | --- | --- | --- | --- | --- |
| Integrated loglik | 153.6 | 5.0 | 0 | 143.6 | 129.06 |
| Penalized loglik | 307.6 | 44.1 | 0 | 219.3 | 91.54 |

**Fixed Effects**

| Predictor | coef | exp(coef) | se(coef) | z | p |
| --- | --- | --- | --- | --- | --- |
| Trial_numberT2 | 1.0276 | 2.7944 | 0.2946 | 3.49 | 0.000485 *** |
| Trial_numberT3 | 1.5255 | 4.5974 | 0.3000 | 5.08 | 3.69e-07 *** |
| Trial_numberT4 | 1.3004 | 3.6708 | 0.2927 | 4.44 | 8.87e-06 *** |
| GenderM | -1.3868 | 0.2499 | 0.6467 | -2.14 | 0.031982 * |

**Significance codes:** *** p < 0.001, ** p < 0.01, * p < 0.05

Supplementary information 4:

**Model Summary**

| Metric | Value |
| --- | --- |
| **Model** | Gamma (log link) |
| **Formula** | Total.Interaction.time..sec. ~ Latency_in_sec + Location + (1 \| Dog_ID) |
| **Data** | data880 |
| **AIC** | 1627.2 |
| **BIC** | 1695.1 |
| **logLik** | -792.6 |
| **Deviance** | 1585.2 |
| **df.resid** | 167 |
| **Dispersion (σ²)** | 0.417 |

**Random Effects**

| Group | Variable | Variance | [Std.Dev](https://std.dev/). |
| --- | --- | --- | --- |
| Dog_ID | Intercept | 0.0665 | 0.2579 |

**Number of obs:** 188, **groups:** Dog_ID (47)

**Fixed Effects**

**Intercept and Continuous Predictor**

| Predictor | Estimate | Std. Error | z value | Pr(>\|z\|) | Signif. |
| --- | --- | --- | --- | --- | --- |
| (Intercept) | 4.0008 | 0.4149 | 9.643 | < 2e-16 | *** |
| Latency_in_sec | 0.0351 | 0.0167 | 2.105 | 0.0353 | * |

**Location (Reference: Chandamari))**

| Predictor | Estimate | Std. Error | z value | Pr(>\|z\|) | Signif. |
| --- | --- | --- | --- | --- | --- |
| LocationGayeshpur | -0.7290 | 0.4811 | -1.515 | 0.1297 |  |
| LocationGhora Gacha | -0.6755 | 0.5138 | -1.315 | 0.1886 |  |
| LocationGokulpur | -0.3670 | 0.5862 | -0.626 | 0.5313 |  |
| LocationJayadebbati | -0.4363 | 0.5867 | -0.744 | 0.4571 |  |
| LocationJonepur | -0.7474 | 0.5870 | -1.273 | 0.2030 |  |
| LocationKalyani A Block | -0.8797 | 0.4370 | -2.013 | 0.0441 | * |
| LocationKalyani B Block | -0.8461 | 0.4323 | -1.957 | 0.0503 | . |
| LocationKalyani Block C | -0.3824 | 0.5078 | -0.753 | 0.4514 |  |
| LocationKalyani D Block | -0.6355 | 0.4807 | -1.322 | 0.1862 |  |
| LocationKalyani ITI More | -1.0540 | 0.5874 | -1.794 | 0.0727 | . |
| LocationKataganj | -0.7095 | 0.4642 | -1.529 | 0.1264 |  |
| LocationKrishnapur | -0.9770 | 0.5923 | -1.649 | 0.0991 | . |
| LocationKulia | -0.3674 | 0.5859 | -0.627 | 0.5307 |  |
| LocationPalladaha | -0.3752 | 0.5883 | -0.638 | 0.5236 |  |
| LocationRina Rd. | -1.8383 | 0.5859 | -3.138 | 0.0017 | ** |
| LocationSiddheswarbati | -0.8075 | 0.5866 | -1.376 | 0.1687 |  |
| LocationSuguna | -0.7655 | 0.5860 | -1.306 | 0.1915 |  |

**Significance codes:** *** p < 0.001, ** p < 0.01, * p < 0.05, . p < 0.1

Supplementary information 5:

Model Summary

| Metric | Value |
| --- | --- |
| **Model** | Gamma (log link) |
| **Formula** | First.lick.to.first.sniff ~ Trial_number + Location + (1 \| Dog_ID) |
| **Data** | data880 |
| **AIC** | 669.0 |
| **BIC** | 743.4 |
| **logLik** | -311.5 |
| **Deviance** | 623.0 |
| **df.resid** | 164 |
| **Dispersion (σ²)** | 0.311 |

Random Effects

| Group | Variable | Variance | [Std.Dev](https://std.dev/). |
| --- | --- | --- | --- |
| Dog_ID | Intercept | 0.1256 | 0.3544 |

**Number of obs:** 188, **groups:** Dog_ID (47)

Fixed Effects

Intercept and Trial Number (Reference: T1)

| Predictor | Estimate | Std. Error | z value | Pr(>\|z\|) | Signif. |
| --- | --- | --- | --- | --- | --- |
| (Intercept) | 1.2835 | 0.4573 | 2.807 | 0.00500 | ** |
| Trial_numberT2 | -0.9255 | 0.1196 | -7.739 | 9.99e-15 | *** |
| Trial_numberT3 | -0.9859 | 0.1188 | -8.298 | < 2e-16 | *** |
| Trial_numberT4 | -0.8536 | 0.1210 | -7.057 | 1.70e-12 | *** |

Location (Reference: Chandamari)

| Predictor | Estimate | Std. Error | z value | Pr(>\|z\|) | Signif. |
| --- | --- | --- | --- | --- | --- |
| LocationGayeshpur | 0.5226 | 0.5258 | 0.994 | 0.3203 |  |
| LocationGhora Gacha | 0.0136 | 0.5549 | 0.024 | 0.9805 |  |
| LocationGokulpur | 1.7938 | 0.6398 | 2.804 | 0.00505 | ** |
| LocationJayadebbati | 0.0563 | 0.6399 | 0.088 | 0.9299 |  |
| LocationJonepur | 0.2822 | 0.6399 | 0.441 | 0.6592 |  |
| LocationKalyani A Block | 0.1284 | 0.4748 | 0.270 | 0.7869 |  |
| LocationKalyani B Block | 0.1603 | 0.4716 | 0.340 | 0.7340 |  |
| LocationKalyani Block C | 0.3608 | 0.5555 | 0.650 | 0.5160 |  |
| LocationKalyani D Block | 0.6219 | 0.5232 | 1.189 | 0.2346 |  |
| LocationKalyani ITI More | -0.4605 | 0.6403 | -0.719 | 0.4720 |  |
| LocationKataganj | -0.1821 | 0.5060 | -0.360 | 0.7190 |  |
| LocationKrishnapur | 0.8320 | 0.6404 | 1.299 | 0.1939 |  |
| LocationKulia | -0.9984 | 0.6397 | -1.561 | 0.1186 |  |
| LocationPalladaha | 0.8750 | 0.6401 | 1.367 | 0.1717 |  |
| LocationRina Rd. | 1.8239 | 0.6400 | 2.850 | 0.00437 | ** |
| LocationSiddheswarbati | -0.7940 | 0.6404 | -1.240 | 0.2150 |  |
| LocationSuguna | 0.7494 | 0.6401 | 1.171 | 0.2417 |  |

**Significance codes:** *** p < 0.001, ** p < 0.01, * p < 0.05, . p < 0.1

Supplementary information 6:

Model summary:

| **Pair Type** | **N** | **Mean Distance** | **SD** | **Median** | **IQR** | **Min** | **Max** |
| --- | --- | --- | --- | --- | --- | --- | --- |
| T1–T2 | 47 | 0.769 | 0.283 | 0.723 | 0.324 | 0.200 | 1.718 |
| T1–T3 | 47 | 0.809 | 0.227 | 0.831 | 0.344 | 0.200 | 1.315 |
| T1–T4 | 47 | 0.779 | 0.236 | 0.760 | 0.260 | 0.000 | 1.286 |
| T2–T3 | 47 | 0.726 | 0.281 | 0.720 | 0.248 | 0.000 | 1.565 |
| T2–T4 | 47 | 0.687 | 0.281 | 0.676 | 0.235 | 0.000 | 1.854 |
| T3–T4 | 47 | 0.691 | 0.298 | 0.638 | 0.299 | 0.000 | 1.357 |

Summary statistics for Optimal Matching (OM) distances across all within-individual trial-pair comparisons.

| **Comparison** | **Estimate** | **SE** | **df** | **t-ratio** | **Adjusted p** |
| --- | --- | --- | --- | --- | --- |
| T1–T3 vs T2–T4 | 48.31 | 12.7 | 230 | 3.798 | 0.0025 |
| T1–T3 vs T3–T4 | 44.91 | 12.7 | 230 | 3.531 | 0.0066 |
| T1–T4 vs T2–T4 | 37.83 | 12.7 | 230 | 2.974 | 0.0378 |

Significant Tukey-adjusted pairwise comparisons from the rank-based mixed-effects model examining OM-distance variation across trial-pair categories (overall model: F = 4.438, p = 0.000704).

| **Metric** | **Value** |
| --- | --- |
| Dogs included | 47 |
| Total pairwise comparisons | 282 |
| Lowest mean OM distance | 0.133 (Dog D28) |
| Highest mean OM distance | 1.231 (Dog D31) |

Overall dataset characteristics and inter-individual variation in behavioural sequence consistency.

Supplementary video 1.

<https://drive.google.com/file/d/1gXmR8mvsMroHc04X9qcb6FZnU3nrNDLi/view?usp=sharing>
